# A novel conserved protein in *Streptococcus agalactiae*, BvaP, is important for vaginal colonization and biofilm formation

**DOI:** 10.1101/2022.03.15.484551

**Authors:** Lamar S. Thomas, Laura C. Cook

## Abstract

*Streptococcus agalactiae* (Group B Strep, GBS) infections in neonates are often fatal and strongly associated with maternal GBS vaginal colonization. Here, we investigated the role of a previously uncharacterized protein, BvaP, in GBS vaginal colonization. BvaP was previously identified as the most highly upregulated gene in the GBS A909 transcriptome when comparing vaginal colonization to growth in liquid culture. We found that expression of BvaP affects GBS adherence to extracellular matrix components and human vaginal epithelial cells and a *ΔbvaP* mutant was significantly decreased in its ability to colonize the murine vaginal tract. Cellular morphological alterations such as changes in cell shape, chain length, and clumping were also observed in a knockout mutant strain. Given its high expression *in vivo*, high degree of conservation among GBS strains, and role in vaginal colonization, BvaP may be an eligible target for GBS vaccination and/or drug therapy.

**IMPORTANCE:** Neonatal GBS disease is a major cause of morbidity and mortality and maternal vaginal colonization is the leading risk factor for disease. Colonization prevention would greatly impact rates of disease transmission, but vaccine development has stalled as capsular polysaccharide vaccines have low immunogenicity *in vivo*. While these vaccines are still in development, addition of a protein conjugate may prove fruitful in increasing immunogenicity and strain coverage across GBS serotypes. Previous research identified *sak_1753* as a gene very highly upregulated gene during murine vaginal colonization. This study reveals that Sak_1753 is required to maintain proper GBS cellular morphology and colonization phenotypes and is required for *in vivo* vaginal colonization in a murine model. We have renamed Sak_1753 Group B Strep vaginal adherence protein (BvaP). The findings of this study indicate that BvaP is important in GBS colonization of the vaginal tract and may be a candidate for vaccine development.

## INTRODUCTION

*Streptococcus agalactiae* (Group B Strep, GBS) colonizes the recto-vaginal tract of approximately 10-35% of the general population (1, 2). GBS is the leading cause of neonatal sepsis worldwide and is accountable for many maternal and fetal bacterial infections, post-infection sequalae, and fetal death (3). GBS may also ascend into the uterus and cause *in utero* infections which may result in premature birth and stillbirth (4). More commonly, GBS is transmitted vertically from a vaginally colonized gravid woman to her child during delivery, likely due to aspiration of contaminated amniotic and bodily fluids, resulting in deadly invasive neonatal diseases. The use of preventive intrapartum antibiotics during labor in GBS-colonized women has been effective against early onset GBS disease but comes with pitfalls including showing no large effect on late onset neonatal GBS diseases (5) and alteration of the neonatal microbiota (6, 7).

The ability of GBS to colonize the vaginal mucosa is essential for pathogenesis of neonatal diseases. While some colonization factors have been described in GBS, many are strain specific or not present in all GBS isolates obtained from colonized women or infected neonates. A more complete understanding of GBS vaginal colonization factors could open new avenues for development of future therapeutics designed to prevent GBS vaginal carriage and neonatal infection without the use of intrapartum antibiotics.

Numerous studies have described examples of GBS interactions with extracellular matrix (ECM) components. The ECM of host mammalian tissue is composed of structural glycoproteins such as fibrinogen and fibronectin, which form a stable macromolecular structure surrounding endothelial and epithelial cells. Researchers proposed that interactions between GBS and ECM components are important for bacterial invasion and host tissue adhesion (8). Described adherence interactions include PilA binding to collagen (9), FbsA and FbsB mediating GBS-fibrinogen binding (8, 10), Lmb binding to laminin (11), and C5a peptidase being involved in fibronectin binding (12). Not all described ECM-adhesins are conserved in all strains of GBS and, as such, the set of surface proteins found in specific GBS strains influences their colonization and virulence potential. Currently, adhesins and other proteins are being explored for vaccine development against pneumonia and sepsis caused by the related organism *Streptococcus pneumoniae* (20) and a similar strategy for GBS vaccination may prove fruitful. In this case, a highly conserved protein adhesin could prove useful as a new therapeutic target (18).

Recent research described the transcriptomic profile of GBS strain A909 growing in liquid culture and compared it to the same strain colonizing the murine vaginal tract for 48-h (21). A hypothetical gene, *sak_1753*, was identified as being most highly upregulated gene in the A909 transcriptome during vaginal colonization. Transcription of this gene was shown to be directly regulated by a two-component system SaeRS. Sak_1753 is conserved in every sequenced GBS strain examined to date although homologs are confined to only a few species of streptococci and enterococci. Our data indicate that Sak_1753 influences mucosal colonization of GBS by altering bacterial biofilm formation ability, attachment to host ECM components, and colonization of the vaginal tract *in vivo*. Deletion of *sak_1753* also results in changes in cell shape, chain length, and clumping ability. Based on our data, we have named Sak_1753 Group B Strep vaginal adherence protein (BvaP).

## MATERIALS AND METHODS

### Bacterial strains, media, plasmids, primers, and growth conditions

All strains and plasmids are shown in Table S1 and the primers used in this study are outlined in Table S2. All GBS strains used in this study were derived from the clinical isolate A909 (22). GBS were routinely grown in Todd-Hewitt medium (BD Bacto) supplemented with 0.2% (wt/vol) yeast extract (Fisher Scientific) (THY) statically at 37°C. Plating was done on THY agar or CHROMagar StrepB agar (DRG). GBS was grown at 37°C unless transformed with a temperature-sensitive plasmid, in which case the growth temperature was 30°C. When necessary, antibiotics were included at the following concentrations for GBS propagation: erythromycin (Erm), 0.5 μg ml^-1^; spectinomycin (Spec), 100 μg ml^-1^; chloramphenicol (Cm), 3 μg ml^-1^. All *Escherichia coli* strains were cultivated in Luria-Bertani (LB) medium or on LB agar. *E. coli* growth was at 37°C unless transformed with a temperature-sensitive plasmid, in which case the growth temperature was 30°C. When necessary, antibiotics were included at the following concentrations for *E. coli* propagation: Erm, 300 μg ml^-1^; Spec, 100 μg ml^-1^, and Cm, 10 μg ml^-1^. Propagation of all shuttle vectors used in this study took place in *E. coli* DH5α, from which plasmid DNA for downstream applications was purified using the Genelute HP Plasmid Miniprep kit (Krackeler) according to the manufacturer’s instructions.

### Creation of mutant and complemented strains

The *ΔbvaP* mutagenesis cassette consists of the spectinomycin resistance gene, *specR*, surrounded by 1000-bp upstream and 996-bp downstream surrounding *bvaP*. The upstream and downstream fragments were PCR amplified from template A909 genomic DNA using primers LC206/LC244 for the upstream region and LC248/LC249 for the downstream region. The *specR* gene was PCR amplified using primers LC246/LC247 from pJC303. The three fragments were cloned into the pMB*sacB* plasmid using 2x Hifi Master Mix (M5520AA; NEB) according to manufacturer’s instructions creating the plasmid pLT001.

Electrocompetent GBS were prepared using the sucrose-free method as previously described (23) with the following modifications: electrocompetent GBS cells were prepared with 1% glycine instead of 2.5% and sucrose concentration was increased to 0.75 M. Transformants were propagated in THY containing Erm and Spec and sucrose sensitivity was confirmed by plating serial dilutions on THY agar with 0.75 M sucrose grown at 30°C. Single crossover intermediates were selected following a temperature shift as previously described followed by passaging in sucrose to select for a double crossover and plasmid excision (23). The final knockout A909Δ*bvaP:*Spec^R^ was sequenced to confirm gene replacement.

For plasmid complementation the *bvaP* gene, including its native promoter were amplified from A909 gDNA by high-fidelity Phusion PCR using LT016/ LT017 primers. The pJC303 plasmid backbone was amplified using primers LT116/LT126 and the chloramphenicol resistant gene *catR* was amplified with LT118/LT119. To construct a strain constitutively expressing *bvaP*, primers LT005/LT009 were used to amplify the pJC303 backbone including a P*recA* promoter and primers LT007/LT008 were used to amplify *bvaP* from A909 gDNA. Gibson assembly for each plasmid was done as described above using 2x Hifi Master Mix creating the complementation plasmid pLT004 (*bvaP_Cm^R^*) and the constitutive plasmid pLT003 (*P_recA_:bvaP_Cm^R^*). Plasmids was electroporated into the A909Δ*bvaP*:Spec^R^ strain. Final constructs were confirmed by sequencing.

### RNA isolation

Following overnight (o/n) growth in THY broth supplemented with the appropriate antibiotics, cultures were diluted 1:20 in fresh THY broth. Once the cells reached an optical density at 600 nm (OD600) of 0.4 to 0.7, 10 mL of cultures were spun down at the same OD, resuspended in 1 mL of RNALater (AM7024; Invitrogen), and incubated at room temperature for 5-10 minutes. For vaginal lavage-induced cells, planktonic cultures were grown as above then pelleted and resuspended in 250 μL of vaginal lavage fluid or PBS as a control. Cells were incubated at 37°C for 1 h then pelleted and resuspended in 1 mL of RNALater and incubated 5-10 minutes at room temperature. Following RNALater treatment, cells were pelleted and stored at −80°C until RNA extraction. RNA was extracted from bacterial pellets using the Ambion RiboPure Bacteria kit (AM1925; Thermo Fisher) according to the manufacturer’s protocol using 7 min of bead beating with a Bead-ruptor 12 (OMNI International) to lyse cells. RNA was DNase I treated using the same kit.

### Preparation of cDNA and qRT-PCR

cDNA preparation was done using the iScript cDNA Synthesis Kit (1708890; BioRad) according to the manufacturer’s instructions, including treatment with RNase H. cDNA was diluted between 1:2 and 1:5, depending on the concentration, and used for qPCR. All primers used in qPCR are listed in Table S2. qRT-PCR was done using the SYBR Green SSo for difficult templates (1725271; Biorad) and a CFX Connect Real Time PCR detection system (788BR01742; Bio-Rad). Gene-specific primers LC060/LC061 (*gyrA*), LT024/LT025 (*bvaP*) were used for amplification. *gyrA*, a housekeeping gene not seen to have differential expression during growth in the vaginal tract, was used as a reference gene. All samples were run in triplicate technical replicates on a single plate, and triplicate biological replicates were used to determine final statistics.

### Growth rate analysis

O/n cultures were diluted 1:20 into fresh THY medium and grown to mid-log phase (OD_600_ = 0.4-0.5). Cultures were diluted down to an OD_600_ of 0.05 in fresh THY medium and 200 μL of each culture was added to a 96-well microplate in 4 technical replicates. Replicates of THY medium only were included as a negative control. The Tecan HP Infinite 200 Pro spectrophotometer was programmed by Tecan i-control software for incubation at 37°C and measurement of OD_600_ every 30 minutes with shaking for 10 secs prior to measurement. Replicates were averaged and plotted as OD_600_ versus time.

### Cell aggregation assay

O/n cultures were diluted 1:10 in 5 mL fresh THY medium and incubated at 37°C until OD_600_ = 0.5. Cells were pelleted by centrifugation then resuspended to a final OD of 0.05 in 50 mL fresh THY medium. Cultures were grown at 37°C until OD = 0.5, vortexed, and placed on the benchtop to settle at room temperature. 1 mL of culture was removed from just below the medium meniscus every 15 minutes to measure OD_600_ using a spectrophotometer (24).

### Biofilm formation assay

O/n cultures were diluted 1:20 into fresh THY and allowed to grow to an OD_600_ value of 0.4. Sterile 9-mm coverslips were added to a 6-well polystyrene microplate which was inoculated with 1 mL of culture. Plates were incubated statically at 37°C with 5% CO_2_ for 24 h to allow cells to attach to coverslip. The medium was aspirated and replaced with fresh THY medium and incubated for another 24 h. To quantify GBS biofilm formation, wells were washed twice to remove cells not bound to the coverslip with 1 mL PBS. Cells were then removed from the wells by incubating with 500 μL of 0.05% Trypsin-EDTA and 0.25% Triton-X100 for 5 min at 37°C. Bacterial counts in each sample were determined by dilution plating on THY agar plates. Quantification was performed in technical triplicate on a single plate and in biological triplicate.

### Scanning electron microscopy

Biofilms grown as described were fixed by flooding the coverslips in fixation solution (2% glutaraldehyde solution, 2% formaldehyde, 150 mM sodium cacodylate buffer, 4% sucrose, 0.15% alcian blue) for 16 h at room temperature. Samples were dehydrated by incubation for 10 min in increasing concentrations of ethanol (50, 70, 80, 95 and 100%) and chemically dried with hexamethyldisilazane (HMDS) o/n in a fume hood (25, 26). The samples were coated with a continuous, conductive, thin-film layer of palladium and platinum alloy prior to their visualization with the Zeiss Supra 55 microscope. SEM images (x50K) were used to measure the length and width of 200 GBS cells using the ImageJ software. Data were analyzed by unpaired t-test using GraphPad Prism 9 software.

### Quantification of GBS A909 chain length

Cultures were grown o/n, and 10 μL of culture was directly placed onto a glass slide with a coverslip and viewed at 1,000x magnification on an Olympus U-LHLEDC. Images were captured of randomly selected visual fields using an attached Olympus DP74 from two separate experiments. At least 100 chains were manually counted from each set of images, for a total of at least 200 chains counted for each strain (27, 28). Data were analyzed by unpaired t-test using GraphPad Prism 9 software.

### Extracellular matrix components adherence assay

Human collagen I (CC050, Millipore Sigma) and human plasma fibronectin (FC010, Millipore Sigma) were diluted to a working concentration of 10 μg/mL in 30% ethanol and added to wells of a 24 well plate. Plates were left to dry o/n in laminar flow hood. Pre-coated human laminin (ECM103, Millipore Sigma) and collagen type IV (ECM105, Millipore Sigma) strip wells were rehydrated by incubating wells with PBS for 15 min at room temperature. O/n cultures of GBS were diluted 1:20 into fresh THY medium and grown to OD_600_ = 0.4-0.5. According to manufacturer’s instructions coated wells were washed once with PBS and 100 μL (strips) or 200 μL (24-well plate) of the bacterial culture was added to each well. Strips and plates were incubated for 4 h at 37°C with 5% CO_2_. Strips and plates were gently washed three times with PBS as above. For quantification, 0.2% crystal violet in 10% ethanol was added and incubated for 5 min at room temperature. Strips and plates were then washed three times with PBS and 200 μL solubilization buffer (50:50 mixture of 0.1M NaH_2_PO_4_, pH 4.5 and 50% ethanol) was added and incubated for 5 min at room temperature. Absorbance was measured at 570 nm on Tecan Infinite PRO spectrophotometer microplate reader.

### VK2 adherence assay

VK2 vaginal epithelial cells were cultured in 24-well tissue culture plates in Keratinocyte Serum-Free Growth Medium (KSFM, Gibco) supplemented with human recombinant epidermal growth factor (rEGF) and bovine pituitary extract (BPE) plus 2% penicillin-streptomycin (Gibco) at 37°C in 5% CO_2_. Cells were grown to a 70-80% confluent state (~10^5^ cells/well) with antibiotics up to 24 h before the assay when the medium was switched to antibiotic-free KSFM + supplements. Immediately before the assay, the eukaryotic cells were washed with PBS. O/n cultures of GBS were diluted 1:20 into fresh THY medium, grown to OD_600_ = 0.4-0.5 (~1×10^8^ CFU/mL), then washed once with PBS, and resuspended in 5 mL KSFM medium. The bacteria were added to the eukaryotic cells at an MOI of ~10:1 and incubated for 1 h at 37°C. Non-adherent bacteria were removed by three washes per well with 1 mL PBS each. Cells were then removed from the wells by incubating with 500 μL of 0.05% Trypsin-EDTA and 0.25% Triton-X100 for 5 min at 37°C with 5% CO_2_. Bacterial counts in each sample were determined by dilution plating on THY agar plates (29). Each adherence experiment was performed in technical triplicate on a single plate and in biological triplicate.

### Mouse model of vaginal colonization

Female outbred CD1 (Charles River) mice aged 6 to 8 weeks were used for all experiments. Experiments were performed as previously described (21, 30). Briefly, 1 day prior to inoculation (day −1), mice were given an intraperitoneal injection of 0.5 mg β-estradiol valerate (Alfa Aesar) suspended in 100 μL filter-sterilized sesame oil (Acros Organics MS) to synchronize estrus. On day 0, mice were vaginally inoculated with bacteria grown to an OD_600_ = 0.4 in 10 μL PBS, at a concentration of ~ 1 × 10^9^ CFU/mL. On days 1, 2, 3, and 5, the vaginal lumen was washed with 50 μL sterile PBS, using a pipette to gently circulate the fluid approximately 6-8 times. The lavage fluid was then collected and placed on ice for no more than 30 min. Vaginal lavage was serially diluted in PBS and plated on CHROMagar StrepB to obtain CFU counts. For long term use, vaginal lavage from healthy uninfected mice were pooled in PBS and frozen at −80°C (21). Murine colonization studies were reviewed and approved by Binghamton University Laboratory Animal Resources (LAR) and by the Binghamton Institutional Animal Care and Use Committee (IACUC) under protocols 803-18 and 857-21.

### Hydrophobicity assay

Two mL of o/n cultures were pelleted by centrifugation at 8,000 x g for 2 min. Pellets were washed twice with PBS then resuspended in 2 mL PBS. 500 μL of o-xylene (Fisher Scientific; AAA11358AP) was added to each sample. Each sample had a negative control consisting only of 2 mL bacteria suspension. Samples were capped and vortexed vigorously for 15 sec. The layers were allowed to separate for 10 min. Hydrophobicity was evaluated through the measurement of the absorbance at OD_600_ of the aqueous fraction divided by its respective control multiplied by 100 then subtracted from 100 (31). The assay was performed in biological triplicate.

### Fractionation of membrane proteins from Gram positive bacteria

An equivalent of 10 mL of culture at OD_600_= 0.6 was pelleted and the supernatant collected. The pellet was resuspended in protoplast buffer (1 M sucrose, 60 mM Tris-HCl, 20 μg/mL lysozyme and 100 U/mL mutanolysin). Following incubation for 45 mins at 37°C, samples were pelleted, and the cell wall fraction (supernatant) moved to a clean microcentrifuge tube. Trichloroacetic acid (TCA) was added to the supernatant samples to give a 10% final solution then incubated on ice for 30 min. Samples were pelleted by centrifugation at maximum speed for 15 min at 4°C. The supernatant was discarded, and the pellet washed once with 500 μL ice-cold acetone and once with 500 μL of ice-cold TCA wash (70% ethanol, bromophenol blue). Pellets were dried using a speed vac centrifuge for 10 min and resuspended in 100 μL 2x Laemmli loading buffer (4% SDS, 20% glycerol, 0.004% bromophenol blue, 0.125M, Tris-Cl, pH 6.8, 10% 2-mercaptoethanol; (32).

The protoplasts (pellet) samples were resuspended in lysis buffer (0.5 M EDTA, 0.1 M NaCl, pH 7.5) and sonicated (85% amplitude, 10s on, 15s off) for 3 min. Samples were centrifuged using a S120-AT2 rotor in the Thermo Scientific Sorvall mTX150 Micro-Ultracentrifuge at 4°C for 1 h at 100,000 x g (33). The supernatant containing the cytosolic fraction was also TCA precipitated and the pellet containing membrane and insoluble fractions was resuspended in 2x Laemmli loading buffer. Fractions were subjected to SDS-PAGE and western blot analysis as described below.

### Western blot analysis

Proteins were run on an SDS-PAGE (11% resolving, 6% stacking), probed with anti-BvaP antibody (Sino Biologicals) and visualized using an ECL kit according to manufacturer’s instructions (80197; Thermo Scientific). The primary antibody was diluted to 0.1 mg/mL and anti-rabbit IgG HRP linked secondary antibody diluted 1:1000 (Cell Signaling Technology; 7074S).

## RESULTS

### BvaP is a novel protein made up of repeated domains conserved within GBS and closely related organisms

BvaP (Sak_1753) from the GBS strain A909 is comprised of 307 amino acids and contains almost five complete repeats of 53 amino acids. Figure 1A depicts the amino acid make-up of BvaP with the proposed signal sequence shown in blue and the 5 repeated domains in black. The putative signal sequence and first repeated domain of BvaP was used as the query sequence in a NCBI database blast to find all possible homologs. BvaP was found in all sequenced GBS strains isolated from several host species including cow and frog. The number of repeats differs between strains but generally ranges from 2 to 6. The sequence is highly conserved within GBS but is also found with less conservation in a few closely related species including *Streptococcus equi* (Fig 1B).

**Figure 1.**
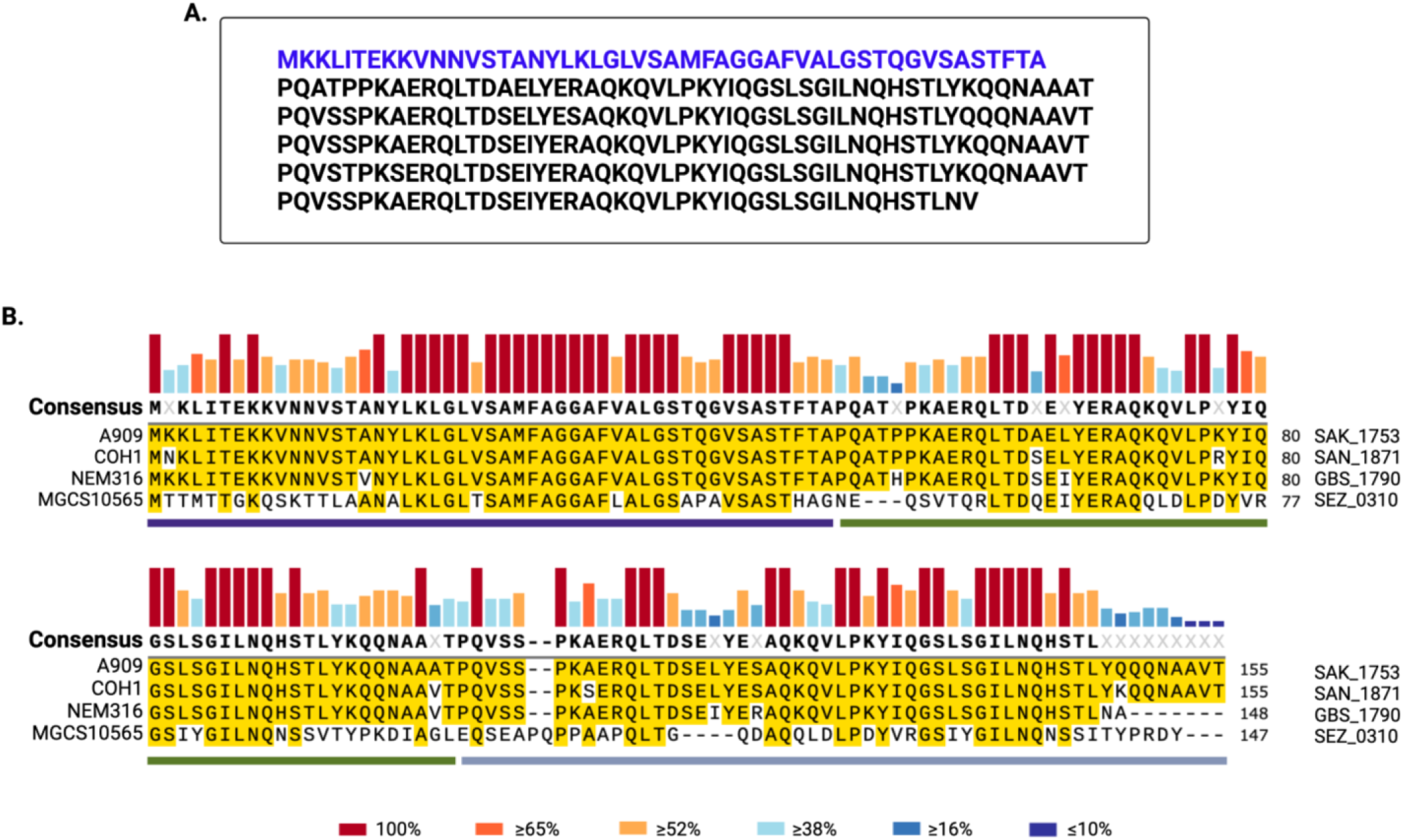
BvaP repeats are highly conserved in all GBS strains. (A) Protein sequence of BvaP (Sak_1753 in strain A909) with putative signal sequence in blue and five repeats in black. (B) The protein sequences of BvaP from three GBS strains (A909, COH1, and NEM316) and one *Streptococcus equi* strain (MGCS10565) were aligned using the Clustal W software in SnapGene. Yellow highlights regions of homology relative to the reference sequence, A909. Vertical bars on top show residue conservation among all strains included. Horizontal bars below represent the putative signal sequence (purple), the first repeated domain (green) and the second repeat (grey).

### Deletion of *bvaP* alters GBS surface-associated phenotypes

Creation of a *bvaP* deletion strain was achieved by allelic exchange in GBS A909 (23). Constitutive expression and complementation strains were also created using plasmids with P*recA* and *PbvaP* promoters respectively in front of *bvaP*. All strains were confirmed by nucleotide sequence analysis and qRT-PCR showed that the constitutive strain expressed *bvaP* approximately 4-fold higher than WT in the same growth conditions (Fig. S1). Growth curves showed no difference in overall growth rate between the mutant and wildtype (WT) strains in THY medium (Fig. 2A), however, a difference in cell clumping was observed in samples incubated without shaking at room temperature (Fig. 2C). Analysis of the sedimentation rate showed significant settling during growth with a decrease in culture turbidity over time for the mutant (Fig. 2B) while the WT turbidity remained relatively constant (Fig. 2B) for longer periods of time. Cell surface hydrophobicity changes have previously been linked to changes in bacterial sedimentation rates (31, 34–37), but we observed no significant difference in surface hydrophobicity between the WT and mutant strains (Fig. S2).

**Figure 2.**
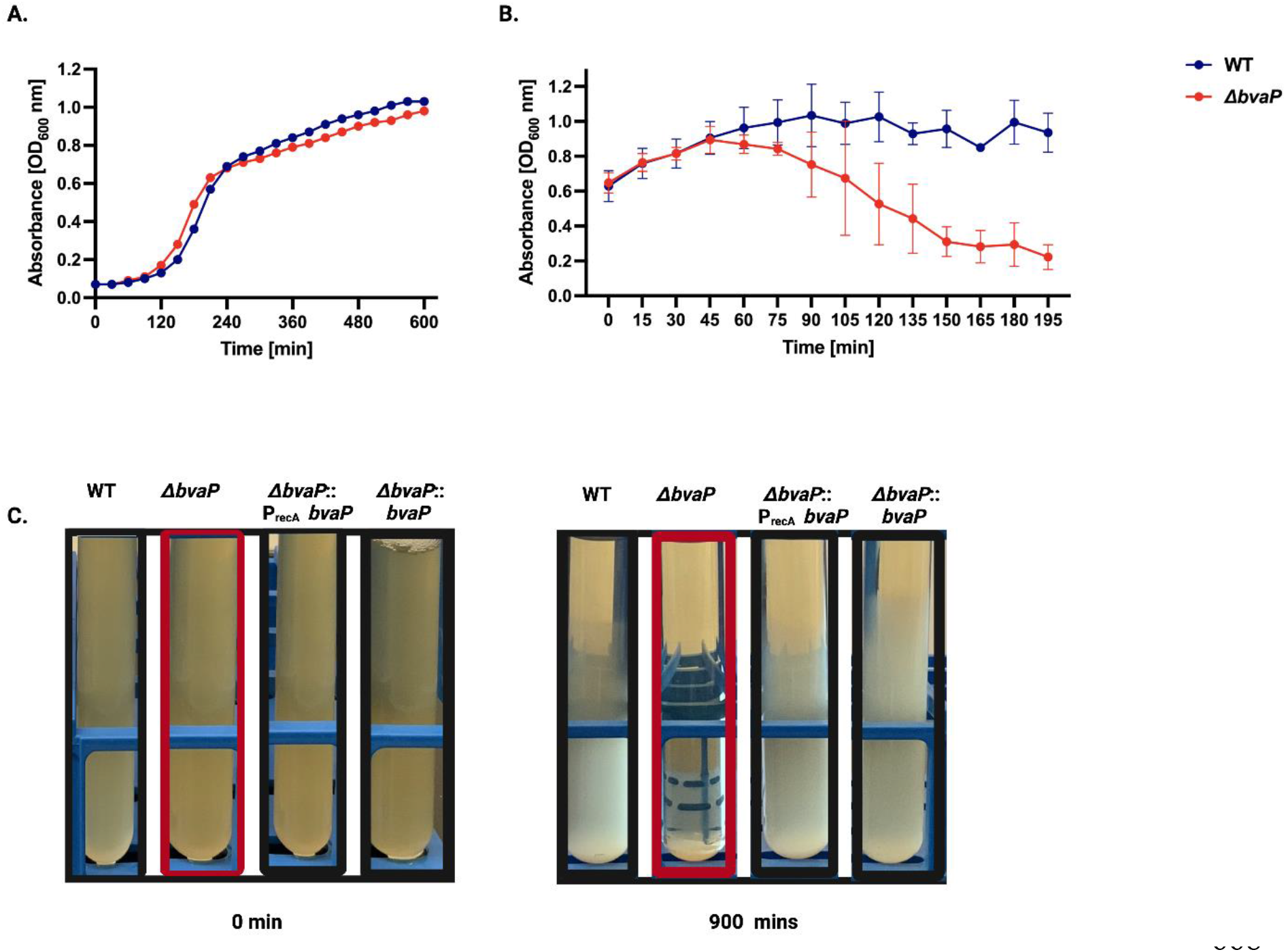
Hyper-aggregation is observed in *A909ΔbvaP* mutant. (A) Growth curve of the WT and A909Δ*bvaP* in THY. Sedimentation curve (B) and macroscopic images (C) of WT and mutant bacterial cells left undisturbed for three hours. Two-way anova was used to for sedimentation rate to analyze statistical significance (p <0.001).

To explore whether the differences in sedimentation rate was linked with cell chaining or clumping, bacteria were grown overnight and stained with crystal violet. A909Δ*bvaP* formed chains of a wider range of size including chains much longer than WT (Fig. 3A). ~200 chains from the WT and mutant strains grown o/n were counted to quantify the number of individual cells in each GBS chain. On average, *ΔbvaP* had significantly longer chains with an average of 20 (±13.9) cells per chain (median value = 14.5), versus the WT which averaged 9 (±6.5) cells per chain (median value = 8.0; Fig. 3B). Complementary, analysis of log phase cells showed the same chain length difference between the WT and mutant strains (not shown).

**Figure 3.**
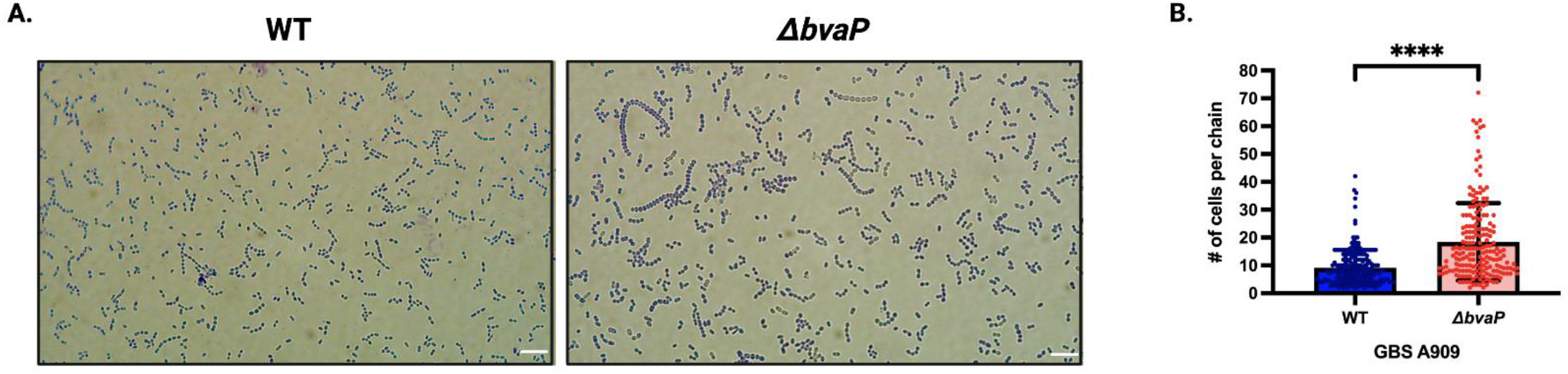
Deletion of *bvaP* increases the average chain length of GBS A909 bacterial cells. (A) Representative images of crystal violet-stained overnight planktonic cultures (scale bar = 10 μm). (B) Average number of cells per chain (n>200). Data was statistically analyzed using Mann Whitney non-parametric T-test; ****, p >0.0001.

Independent of chain length, a difference in cell size and morphology between the two strains was observed microscopically. To evaluate the morphological differences between the WT and mutant strains, scanning electron micrographs were obtained to visualize the details of the bacteria surface morphology and to measure cell length and diameter. Interestingly, the length and width of the mutant cells were observed to be significantly different than that of WT cells. WT cells appeared more ovoid with an average length of 0.80 (±0.10) and a width of 0.70 μm (±0.06) while the mutant cells were more spherical, the length was shorter at 0.72 μm (±0.10) and width slightly wider at 0.72 μm (±0.08; Fig. 4A and B). SEM also exposed variation in the division septa. In the absence of BvaP, the septum is observably narrower than the WT in some cells (Fig. 4C).

**Figure 4.**
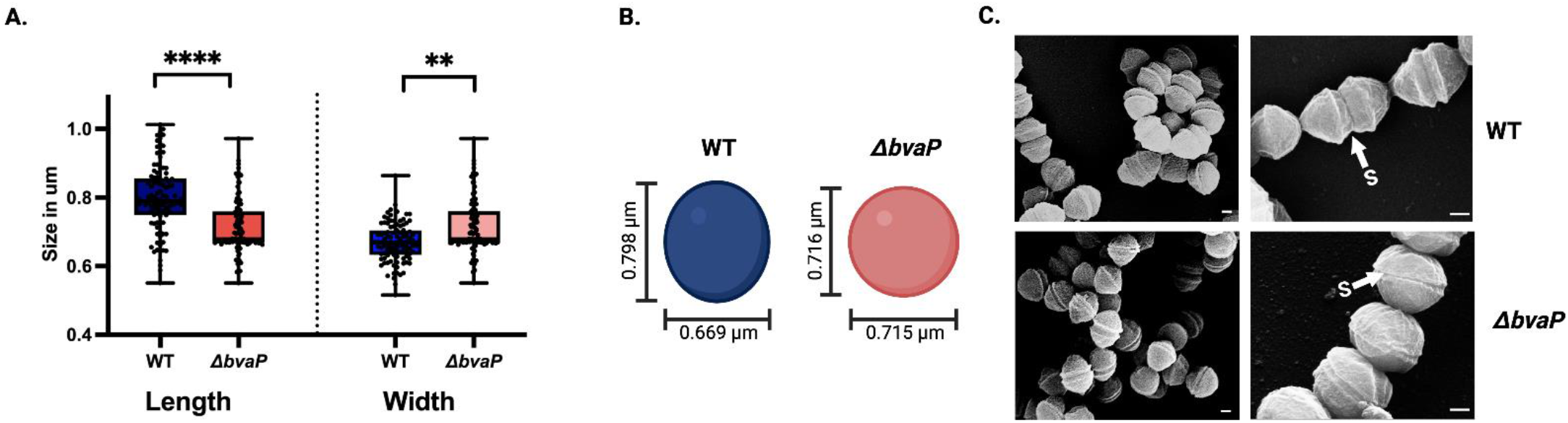
Deletion of *bvaP* caused modifications to GBS cell shape and morphology. (A) Cell size of log phase GBS planktonic cells. (B) Diagrammatic illustration of WT A909 cell diameter compared to ΔbvaP. (C) Representative scanning electron micrograph of WT and mutant GBS cells coated with palladium and titanium for surface morphology imaging. S represents the diving septum; scale bar from left to right = 0.2 μm. p value determined by Mann-Whitney U test; **, p <0.01; ****, p <0.0001.

### BvaP is localized to the cell wall and membrane with a portion secreted or released

To determine the subcellular localization of BvaP, western blot analysis was employed. Polyclonal anti-BvaP antibody was generated by Sino Biologicals against two peptides from BvaP. A cellular differential fractionation protocol was used to separate the cytosolic and membrane fractions. Protein from culture supernatant, and cell wall were separated and assessed for BvaP. Under laboratory growth conditions, BvaP is found secreted in the culture supernatant and localized in the cell wall fraction of the cell (Fig. 5).

**Figure 5.**
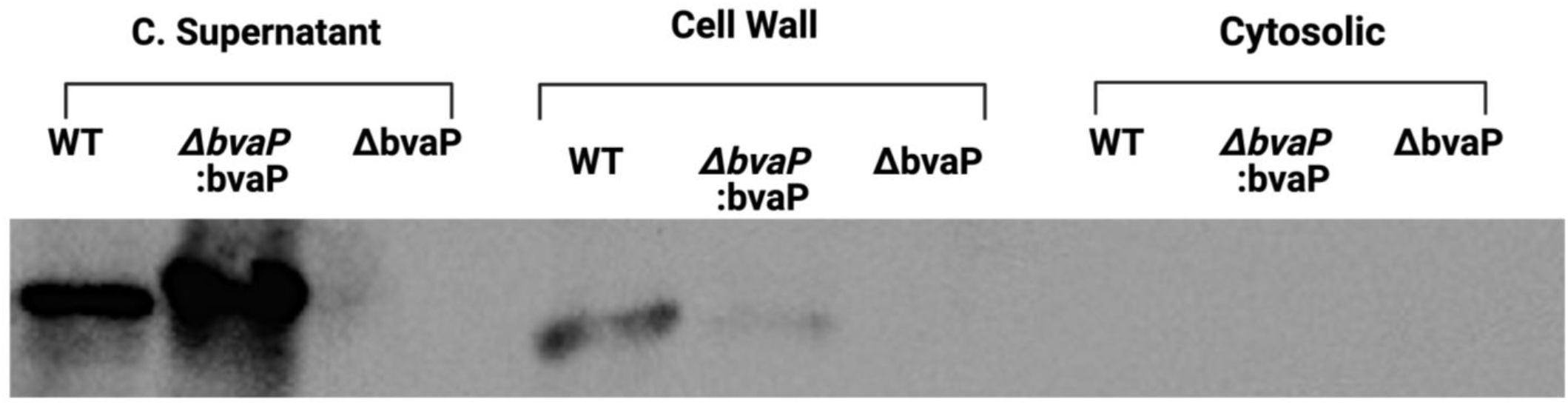
BvaP is found in the cell supernatant and cell wall compartments. Western blot analysis of whole cell lysates fractioned by ultracentrifugation to identify the subcellular localization of BvaP. Samples were loaded on SDS-PAGE with loading buffer containing 2% β-mercaptoethanol, heated at 95°C for 10 min, anti-BvaP antibody (Sino Biological, rabbit polyclonal, 0.1mg/mL with 4% BSA in T/TBS) at 4°C overnight followed by anti-rabbit IgG-HRP conjugated (1,000-fold diluted in T/TBS) at room temperature for 2 h.

### BvaP aids in GBS adherence to collagen I and human vaginal epithelial cells (VK2)

GBS adherence to host cells is an important preliminary step for successful mucosal colonization. To ascertain the role of BvaP in the colonization of the host mucosa, we examined adhesion of the WT and mutant strains to host extracellular matrix components (ECMs) and a human vagina epithelial cell line, VK2. To examine if BvaP interacts with human ECM components; fibrinogen, fibronectin, laminin, plasminogen, and collagens I and IV were selected for adherence assays. Our data demonstrate that deletion of *bvaP* does not affect binding of A909 to collagen IV, plasminogen, and laminin. However, constitutive expression of *bvaP* resulted in a significant increase in adherence to human collagen I while deletion of *bvaP* significantly reduced adherence (Fig. 6A). Overexpression of *bvaP* also resulted in significantly increased binding of bacteria to fibronectin and fibrinogen. Published RNA-seq data showed that expression of *bvaP* is low in liquid culture (21) so comparison of the knockout and constitutive expression strains may be more physiologically relevant in laboratory medium.

**Figure 6.**
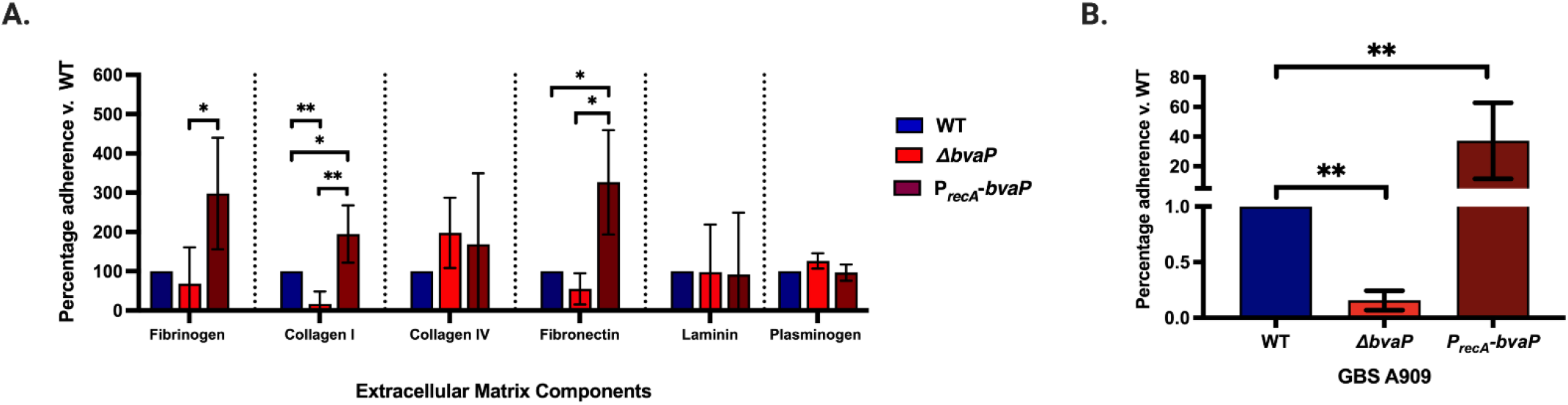
BvaP is necessary for proficient binding to some ECMs and human vaginal epithelial cells *in vitro*. (A) Adhesion assays with extracellular matrix components: human fibrinogen, collagens I and IV, fibronectin, and laminin. Significant binding highlighted in brown. Assays were completed in at least technical and biological triplicate. Statistical significance was analyzed by -parametric Mann-Whitney test for the ECM components (n = 4), *p < 0.05, **p <0.01. Data represent the mean (± standard deviation, S.D.). (B) Adhesion assay with human vaginal epithelial cells (VK2) at MOI of 10 shows decreased binding of the *bvaP* mutant strain and increased binding for the strain constitutively expressing *bvaP* compared to WT. Statistical significance was analyzed by one-way ANOVA (n = 3). **, p < 0.01. Data represent the mean (± S.D.).

For cell culture adherence assays, VK2 cells were grown in 24 well plates to ~70-80% confluence. Bacterial cells were grown to log phase (OD_600_ = 0.4-0.5), washed with PBS, and resuspended in antibiotic-free KSFM at a multiplicity of infection (MOI) of ~10. Following a 30-minute incubation with bacteria, the VK2 cells were washed to remove non-adherent bacteria and the adherent cells were quantified. Constitutive expression of *bvaP* using the *P_recA_-bvaP* plasmid resulted in a significant increase in GBS adherence while deletion of *bvaP* resulted in significantly decreased binding of GBS to VK2 cells (Fig. 6B).

### *In vitro* biofilm formation by GBS is inhibited in the absence of *bvaP*

The formation of bacterial biofilms is often proposed to be important for colonization of the host mucosa. To examine the impact of BvaP on *in vitro* GBS biofilm formation, 24- and 48-hour biofilms were grown on glass coverslips in 6-well plates. Biofilms were examined using both SEM and live/dead staining in conjunction with fluorescent microscopy to visualize biofilm structure. The *ΔbvaP* mutant was unable to form the structurally intricate biofilms seen in the WT samples (Fig. 7A) and the biofilm that was formed consisted mostly of cells that stained red with a Live/Dead stain indicating increased membrane permeability (Fig. 7B). To check the cell viability during biofilm growth, biofilms were grown as previously described. A solution of 0.05% trypsin -EDTA and 0.25%Triton X-100 was used to detach biofilm from coverslip and bacteria was quantified through plate dilution and CFU counts. Mutant biofilm counts were lower than WT at 24-h but not significantly whereas there was a statistically significant difference at 48-h (Fig. S3).

**Figure 7.**
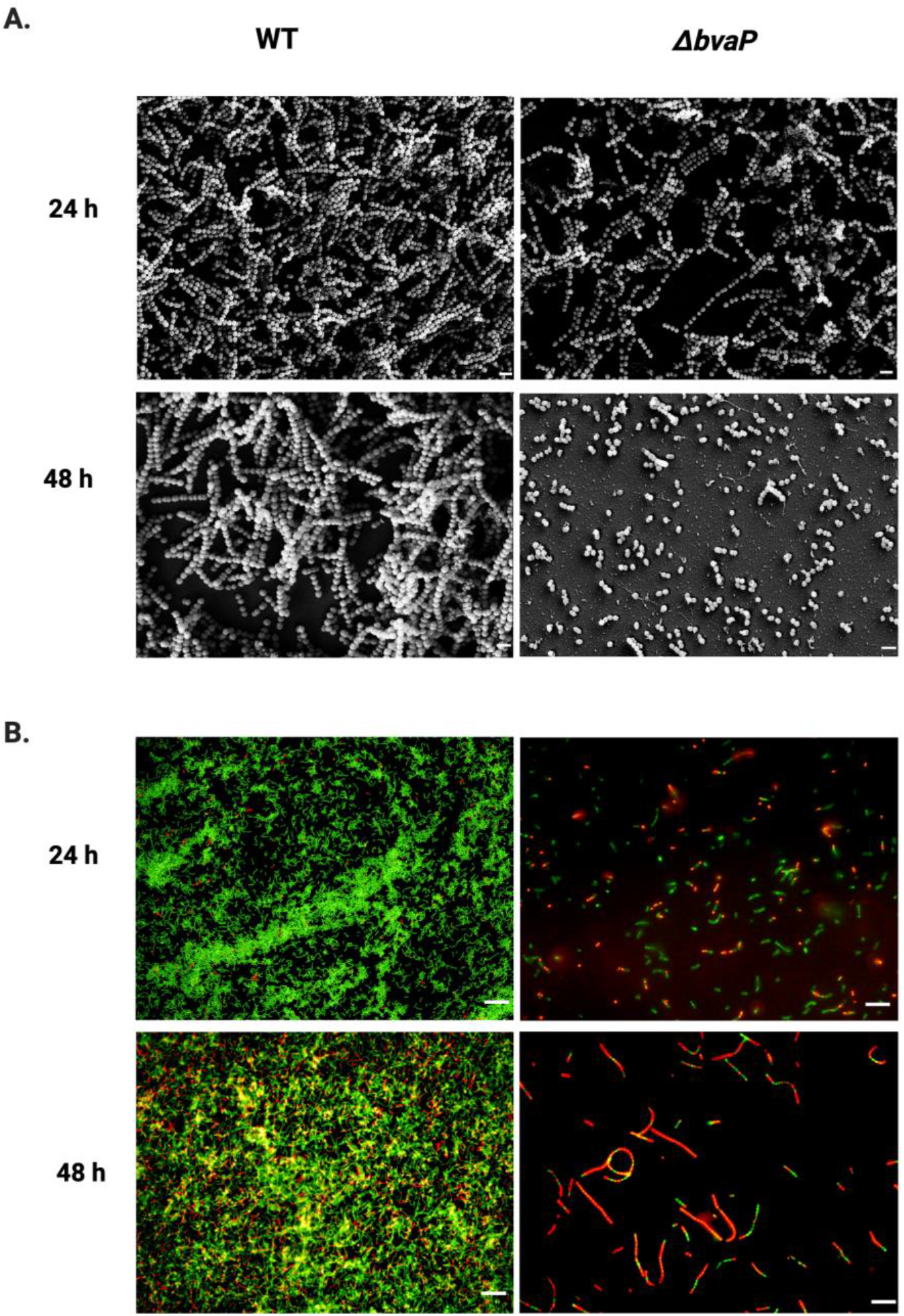
BvaP is required for efficient biofilm formation on glass surfaces with and without vaginal lavage induction. (A) Scanning electron micrograph of GBS WT and mutant □bvaP strains. Scale = 2 μm (B) BacLight Live/Dead stain (green-SYTO9 and red- propidium iodide) of GBS biofilm. A909 biofilm formation with mice vagina lavage induction (pink arrow indicate microcolonies formed in response to vagina lavage). Scale = 10 μm. (C) BacLight Live/Dead stain of GBS log phase planktonic cells. Scale = 10 μm.

### BvaP is required for efficient *in vivo* murine vaginal colonization

To evaluate the role of BvaP in the GBS colonization of the vaginal tract, a murine vaginal colonization model was employed. Mice were injected intraperitoneally with β-estradiol to synchronize their estrus cycles on day −1. On day 0, mice were inoculated intravaginally with ~10^7^ CFU of bacteria in 10 μL. On subsequent days, vaginal washing was done followed by plating on GBS selective agar to quantify bacterial numbers. Compared to WT, *ΔbvaP* showed significantly less GBS colonization on days 1-3 and a trend toward decreased colonization on day 5 indicating an important role for this protein in vaginal adherence, especially early on in the mucosal colonization process (Fig. 8).

**Figure 8.**
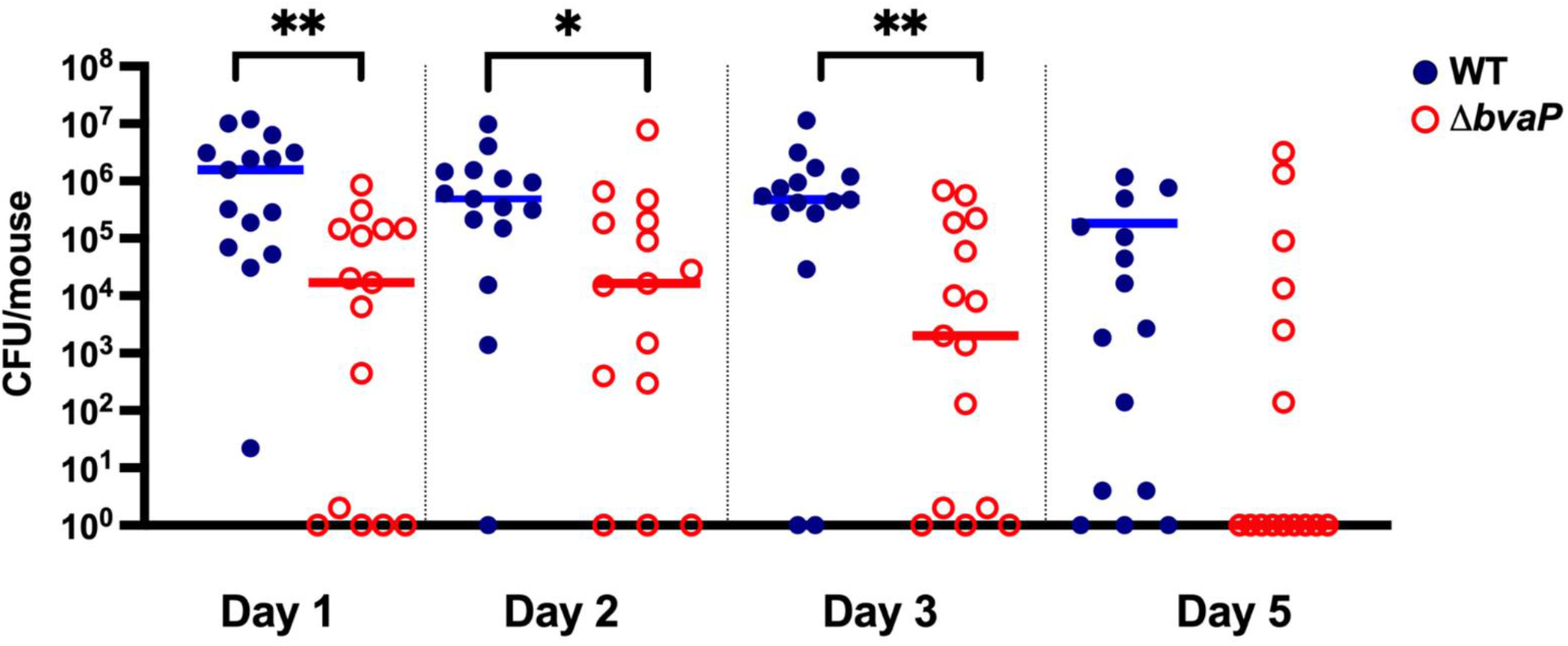
BvaP is required for in vivo murine vaginal colonization. Estradiol synchronized CD 1 mice were colonized with 10^7^ CFU/mouse of WT or *ΔbvaP* strains. Bar is set at the median value. Data were analyzed using Mann-Whitney test; *, p <0.005; **, p < 0.001.

## DISCUSSION

A previously published transcriptomic study identified an uncharacterized gene, *sak_1753*, as the most highly upregulated gene in GBS strain A909 when comparing vaginal colonization to growth in laboratory culture (21). Here, we have begun to characterize Sak_1753, now identified as Group B Strep vaginal adherence protein (BvaP). While BvaP is found in all sequenced strains of GBS, few homologs are present in even closely related species such as *Streptococcus pyogenes* and there are no predicted protein family or domain homologs using protein prediction software Pfam and Phyre. Here, we present the first information on the effects of BvaP on GBS surface morphology and phenotypes associated with vaginal colonization.

Initial observations indicate that deletion of *bvaP* from the A909 genome alters the ability of cells to aggregate in laboratory medium (Fig. 2) and changes the overall cell shape resulting in bacteria that are significantly wider than WT (Fig. 4C). Additionally, the chains formed by *ΔbvaP* mutants are significantly longer than WT (Fig. 3), potentially denoting incomplete cell division. Bacterial cell shape and chain length changes in particular mutants or growth conditions have been reported for many chain-forming bacteria. For streptococci, the regulation of chain length has been reported to be dependent on cell wall-associated autolytic activity and the presence of autolysins (38, 39) as well as environmental factors such as pH (38), growth medium (40), and salt concentration (39), among others. Autolysins are important for bacteria cell-to-host surface adhesion, cell division (41), antibiotic resistance, peptidoglycan turnover and spore development (42). Peptidoglycan breaks are both naturally occurring and regulated by autolysins. As such, a deletion in one autolysin gene can keep large peptidoglycan molecules intact leading cells to stay together after a cell division cycle. Mercier et al. 2002, reported the formation of long chains in an *Lactococcus lactis* strain with a mutation in the peptidoglycan hydrolase, *acmA*. They highlighted that this increase in chain length reflects a reduction in the frequency of peptidoglycan breaks and they observed reduced adherence to solid surfaces as well as reduced biofilm formation in a mutant with increased chain length. Other autolysins such as LytR/S (42, 43) and LrgA (44) in *S. aureus*, LytB in *S. mutans* (45) and *S. pneumoniae* (46), LytA in *S. pneumoniae* (47),, and AtlA (49) in *Enterococcus faecalis*, have been reported to cause similar changes in chain length and morphology. While BvaP has no known homology to autolysins, the phenotypic similarities are intriguing and could signify that *bvaP* plays a role in peptidoglycan formation or breakage at the cell surface, potentially through structural influence.

There are also studies linking chain length with pathogenesis, showing that long chain cells may promote adherence and colonization due to an increase in surface area which allows for multivalent adhesive interactions (39), while short chains are associated with more invasive diseases such as meningitis (40). Alternatively, other studies show longer chain phenotypes in mutants with decreased adherence and colonization ability like that seen in our studies with BvaP (50, 51). In addition, Li et al., described an increase in *S. pneumoniae* chain length in a mutant strain correlated with an increase in cell death through C3 mediated neutrophil opsonophagocytosis (52). The reduced ability to form biofilms is also often linked to phenotypes such as increased aggregation and chain length (31). Deletion of *bvaP* was correlated with a decrease in biofilm formation (Fig. 7). Biofilm formation by *ΔbvaP* at 24-h was markedly reduced compared to WT and by 48-h there is a noticeable dearth of biofilm compared to the WT.

Cell surface hydrophobicity is an important element for bacterial surface attachment. Alterations in surface hydrophobicity have been associated with changes in biofilm formation (53), sedimentation and phenotypes (34, 35). We found no change in cell surface hydrophobicity in the *ΔbvaP* mutant, suggesting that the overall cell surface chemistry remains unchanged in the absence of BvaP (Fig. S2). In addition, capsular alterations may be associated with aggregative phenotypes (54) but we did not observe differences in capsule between our WT and mutant cells.

BvaP was not detected in the bacteria cytosol but was found in both on the cell surface and in the extracellular space. Detection of BvaP in the cell suface (Fig. 5) suggests that BvaP is potentially anchored in the cell wall or cell membrane. In Gram positive bacteria, proteins may be covalently anchored to the cell wall through a highly conserved carboxy-terminal motif Leu-Pro-X-Thr-Gly (LPXTG) which is cleaved and then attached to an amino group of the processed protein in the peptidoglycan via sortase (55–57). Less commonly, anchorage is through lipoproteins where proteins are tethered on an amino-terminal lipid-modified cysteine (58, 59). The mechanism by which BvaP, which lacks a canonical LPXTG motif, is tethered to the surface or released extracellularly, either actively or passively, has yet to be explored.

Binding and colonization of the host mucosa often involves interactions between bacteria and the host extracellular matrix (ECM) components. Proteins expressed by streptococci can mediate interactions with host ECM components to contribute to its adherence, colonization, invasion, and evasion from host defenses (60, 61). Collagen is the most prevalent ECM component in humans and plays a critical role in maintaining the integrity of most tissues (62). Previous studies have shown conflicting results as to whether GBS binds collagen I to mediate host attachment. Dramsi et al. argued that that GBS strain A909 does not bind to collagen I but instead binds to fibrinogen to mediate attachment to the host (9) yet Avilés-Reyes et. al. (63), identified significant binding to collagen I. Experiments in the former study were conducted using rat tail collagen I compared to latter study which used human collagen, which may explain the difference in findings. In this study, we found that GBS A909 binding to human collagen I is at least partially mediated by BvaP (Fig. 6A). We observed GBS binding to both fibrinogen and fibronectin (Fig. 6A), corroborating previous findings(8) and these interactions also appear to be affected by the presence of BvaP. In addition to specific ECM components, we also undertook adherence assays using immortalized cultured vaginal epithelial VK2 cells. Cells in which *bvaP* was deleted had greatly decreased ability to bind VK2 cells and this adherence defect could be complemented by addition of *bvaP in trans* (Fig. 6B).

Results of our murine vaginal colonization assays agree with our *in vitro* data indicating a role for BvaP in host binding. Mice inoculated with *ΔbvaP* were colonized significantly less over the 5-day period than mice inoculated with the WT. Since our data indicate that BvaP is important for binding some ECM components, it would be interesting to examine if the adherence ability of BvaP to the host vaginal mucosa is dependent on interactions with these ECMs specifically or to overall changes in cell surface properties affecting colonization more generally.

Collectively, our data provides important initial characterization of a previously hypothetical protein, Sak_1753, now BvaP, conserved throughout GBS strains. BvaP is integral to maintaining GBS cell morphology, chain length, and *in vitro* biofilm formation ability, and is required for efficient adherence to vaginal epithelial cells, both in *in vitro* adherence assays and *in vivo* murine vaginal colonization experiments. It is possible that these adherence properties are mediated by ECM components such as collagen I, fibrinogen, and fibronectin although future studies are needed to determine the role of these interactions *in vivo*. Cumulative data from this and previous studies are combined to create a model of colonization shown in Figure 9. The expression of *bvaP* is regulated by the SaeRS two component system which is initiated upon the reception of a signal from the host vaginal environment (21). The downstream effect of this signaling is translation of BvaP, resulting in increased adhesion to host vaginal mucosa, helping to initiate and maintain colonization (Fig. 9). More research will be needed to determine whether the colonization defect observed in the *ΔbvaP* mutant is related specifically to chain length, cell surface property alterations, host immune interactions, specific adhesion binding or some combination of these.

**Figure 9.**
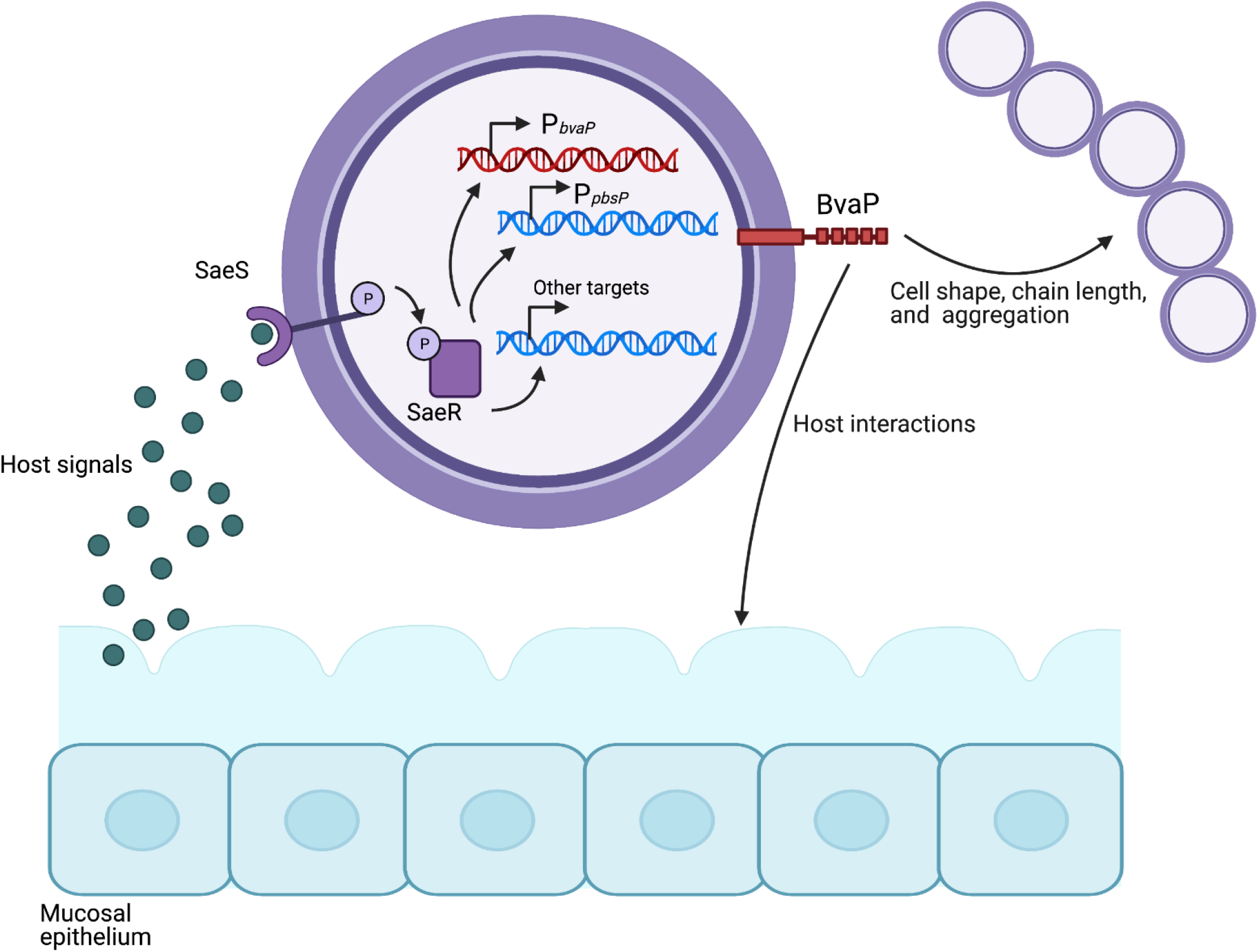
Working model of BvaP regulation by SaeRS two component system during vaginal colonization. The system is initiated when the histidine kinase, SaeS, senses a signal in the vaginal mucosa stimulating autophosphorylation of SaeS and phosphorylation of the response regulator, SaeR. SaeR directly upregulates expression of *bvaP*, resulting in increased biofilm formation and binding to the host mucosa.

## Acknowledgements

We thank Jennifer Amey (S3IP – Small Scale Systems Integration and Packaging, Binghamton University, Binghamton, New York, USA) for her technical assistance in capturing SEM images. Some figures were created using Biorender.com

